# FOXO mediates organismal hypoxia tolerance by regulating NF-κB in *Drosophila*

**DOI:** 10.1101/679605

**Authors:** Elizabeth C Barretto, Danielle M Polan, Amy N Beever-Potts, Byoungchun Lee, Savraj S Grewal

## Abstract

Exposure of tissues and organs to low oxygen (hypoxia) occurs in both physiological and pathological conditions in animals. Under these conditions, organisms have to adapt their physiology to ensure proper functioning and survival. Here we define a role for the transcription factor FOXO as a mediator of hypoxia tolerance in *Drosophila*. We find that upon hypoxia exposure, FOXO transcriptional activity is rapidly induced in both larvae and adults. Moreover, we see that *foxo* mutant animals show misregulated glucose metabolism in low oxygen and subsequently exhibit reduced hypoxia survival. We identify the innate immune transcription factor, NF-KappaB/Relish, as a key FOXO target in the control of hypoxia tolerance. We find that expression of Relish and its target genes are increase in a FOXO-dependent manner in hypoxia, and that *relish* mutant animals show reduced survival in hypoxia. Together, these data indicate that FOXO is a hypoxia inducible factor that mediates tolerance to low oxygen by inducing immune-like responses.

## INTRODUCTION

Oxygen is essential for normal growth, development and functioning of tissues and organs. However, while the air we breathe contains ~20% oxygen, even under healthy physiological conditions, our cells and tissues receive considerably lower levels. These can be anywhere from 1 to 10% oxygen depending on the tissue (McKeown, 2014). Hence, our tissues and organs need to function and maintain homeostasis at low levels of oxygen. This aspect of normal physiology is often neglected in tissue culture experiments where cells are routinely maintained in 20% oxygen. In addition, many diseases such as heart disease, stroke and chronic lung disease are characterized by severe oxygen deprivation (hypoxia) (Semenza, 2011). This hypoxia has deleterious effects on tissue metabolism and function, and can lead to death. Understanding how cells, tissues and organisms adapt to low oxygen is therefore an important question in biology.

One central hypoxic mechanism involves induction of the HIF-1α transcription factor, which can control the expression of a diverse array of target genes that maintain cellular homeostasis in low oxygen (Semenza, 2014). The importance of HIF-1α has been shown by loss of function genetic analysis in model organisms such as *C elegans*, *Drosophila* and mice. For example, in *C elegans* and *Drosophila*, which are normally quite hypoxia-tolerant, HIF-1α mutants die when exposed to low oxygen (Centanin et al., 2005; Jiang et al., 2001; Li et al., 2013). Tissue-specific mouse knockouts have also shown how HIF-1α can control organ-level and whole-body adaptation to low oxygen in both physiological and pathological conditions (Boutin et al., 2008; Cramer et al., 2003; Huang et al., 2004; Mason et al., 2004; Schipani et al., 2001; Tomita et al., 2003). Compared to our understanding of HIF-1α, however, less is known about other transcription factors that are important in mediating hypoxia adaptation in animals.

The conserved transcription factor Forkhead Box-O (FOXO) is an important mediator of adaptation to stress in animals (Webb and Brunet, 2014). Studies in *Drosophila* have provided important insights into the role of FOXO as a regulator of organismal physiology. Here, different environmental stressors, such as starvation, oxidative stress, pathogens and ionizing radiation, have been shown to induce FOXO transcriptional activity (Borch Jensen et al., 2017; Dionne et al., 2006; Junger et al., 2003; Karpac et al., 2009; Karpac et al., 2011). Once induced, FOXO then directly controls the expression of an array metabolic and regulatory genes that together function to maintain organismal homeostasis and survival (Alic et al., 2011; Birnbaum et al., 2019; Gershman et al., 2007; Teleman et al., 2008). Indeed, genetic upregulation of FOXO is sufficient to promote stress resistance in *Drosophila*, and it is one of the most effective ways to extend lifespan (Alic et al., 2014; Demontis and Perrimon, 2010; Giannakou et al., 2004; Hwangbo et al., 2004; Kramer et al., 2008).

In this paper, we report our work using *Drosophila* to explore hypoxia tolerance. In their natural ecology, *Drosophila* grow in rotting, fermenting food rich in microorganisms – an environment likely characterized by low ambient oxygen (Callier et al., 2015; Harrison et al., 2018; Markow, 2015). Probably as a consequence of this, they have evolved mechanisms to tolerate hypoxia (Centanin et al., 2008; Lee et al., 2019; Li et al., 2013). Here we show that induction of FOXO is one such mechanism and that it functions by regulating an immune-like response.

## RESULTS

### Hypoxia induces FOXO activity

The main way that FOXO is regulated is through nuclear-cytoplasmic shuttling. In order to determine if hypoxia exposure could induce FOXO, we transferred third instar larvae growing on food to either moderate (5% oxygen) or severe hypoxic environments (1% oxygen) and then stained for FOXO localization using an anti-FOXO antibody (Figure 1A). We saw that exposure to hypoxia caused FOXO relocalization from the cytoplasm to the nucleus of fat body cells (Figure 1A). This effect was rapid; nuclear relocalization occured within 15 minutes of exposing larvae to hypoxia (Figure S1A). We next examined the effects of hypoxia on the expression of *4e-bp*, a well-characterized FOXO target gene. We measured mRNA levels of *4e-bp* using qRT-PCR in whole third-instar larvae exposed to either 5% or 1% oxygen. We saw that *4e-bp* levels were strongly increased in control (*w*^*1118*^) larvae exposed to both hypoxic conditions (Figure 1B, C). As with the FOXO nuclear localization, this increase in *4e-bp* was rapid and was seen within 15-30 minutes following hypoxia exposure (Fig S1B). However, the hypoxia-induced increase in *4e-bp* mRNA levels was largely abolished in *foxoΔ94*, a deletion line that is a null mutant for the *foxo* gene (Slack et al., 2011)(Figure 1B, C). We also examined the effects of hypoxia in adults. We exposed adult females to 1% O_2_ and found that, as in larvae, *4e-bp* levels were increased in control (*w*^*1118*^) animals and that this effect was blunted in *foxo* mutants (Figure 1D). Finally, we examined the tissue pattern of *4e-bp* induction by examining LacZ staining in *thor-LacZ* flies, which is a LacZ-enhancer trap in the *4e-bp* gene locus (Bernal and Kimbrell, 2000). We found that larvae exposed to 2 hours of 5% O_2_ showed increased LacZ staining in the majority of larval tissues including the fat body, the intestine, and the body wall muscle (Figure 1E), suggesting that the hypoxia induction of FOXO activity is not tissue-restricted. Together, these data indicate that exposure to hypoxia in both *Drosophila* larvae and adults results in rapid induction of FOXO transcriptional activity.

**Figure 1.**
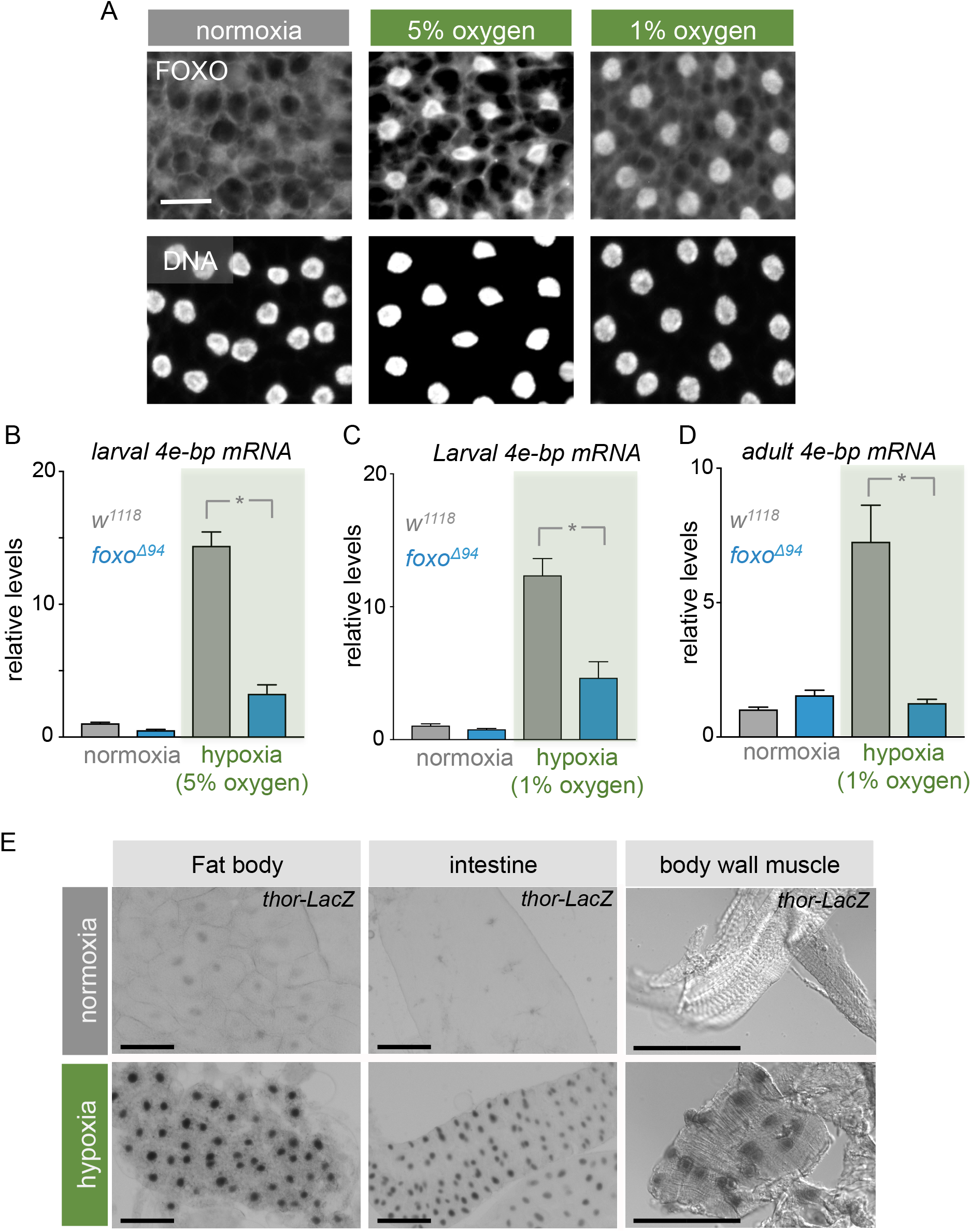
Hypoxia induces FOXO activity. (A) FOXO staining of 96-hour AEL *w*^*1118*^ larval fat bodies following exposure to hypoxia for two hours. Nuclei are stained with Hoechst (bottom panels). Scale bar is 25 *μ*m. (B) *4e-bp* mRNA levels measured by qRT-PCR in control (*w*^*1118*^) and *foxo* mutant (*foxoΔ94)* following B) 6 hours of 5% O_2_ hypoxia in larvae, C) 6 hours of 1% O_2_ hypoxia in larvae, or D) 16 hours of 1% O_2_ hypoxia in adults. N>6 cohorts of animals per condition. Data represent mean + SEM. * p,0.05, two-way ANOVA followed by post-hoc t-test. (E) LacZ staining in tissues of thor-LacZ larvae following two-hour exposure to 5% O_2_. Scale bar is 100μm

### FOXO is required for hypoxia tolerance

Is FOXO activation required for *Drosophila* survival in low oxygen? To find out, we measured hypoxia survival in *foxoΔ94* animals. Under standard laboratory conditions (rich food, normoxia) *foxo* mutant animals are viable (Slack et al., 2011). We therefore examined how well these mutants tolerate low oxygen. We first examined hypoxia in larvae. Control (*w*^*1118*^) and *foxo* mutant embryos were allowed to develop in normoxia and then newly hatched larvae were transferred to hypoxia (5% oxygen) for the duration of their larval period, before being returned to normoxia. We then counted the number of animals that developed to viable adults. We found that the *foxo* mutant animals reared in hypoxia had a significant decrease in viability compared to control animals (Figure 2A). We next examined hypoxia survival in adults. Control *(w^1118^)* and *foxo* mutant animals were exposed to either severe hypoxia (1% oxygen) for 24 hours or anoxia (0% oxygen) or 6 hours. After these low oxygen exposures, flies were returned to normoxia and the number of surviving animals counted. As observed in larvae, we found that the adult *foxo* mutant animals showed significantly deceased survival in both the hypoxic and anoxic conditions (Figure 2B, C). During severe hypoxia and anoxia, adult flies become immobile. However, when *foxo* adults were exposed to starvation instead of hypoxia for 24 hours, there was no effect on viability, indicating that the decrease in hypoxia survival in *foxo* mutants is not simply a consequence of reduced nutrient intake as a result of immobility (Fig S2). Together, our data indicate that FOXO activation is required for organismal survival in low oxygen in both developing larva and adults.

**Figure 2.**
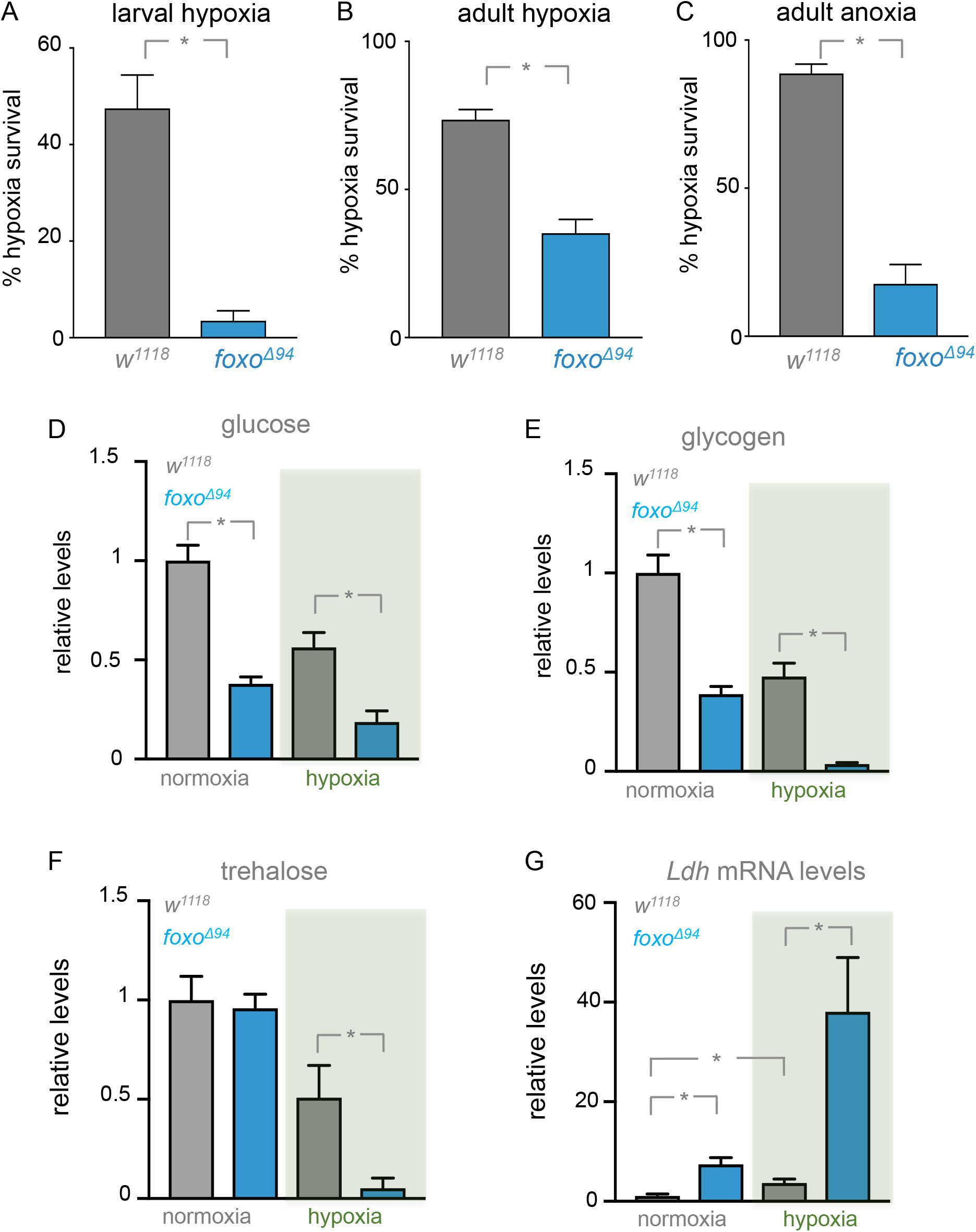
FOXO is required for hypoxia tolerance. (A) Control (*w*^*1118*^) and *foxo* mutant (*foxoΔ94*) animals were exposed to hypoxia (5% O_2_) throughout their larval period, before being returned to normoxia as pupae. The percentage of flies that eclosed as viable adults were then counted. (B,C) Adult control (*w*^*1118*^) or *foxo* mutant (*foxoΔ*^*94*^) flies were exposed to either, B) 24 hours of 1% O_2_ or C) 6 hours of 0% O_2_, before being returned to normoxia. The percentage of viable flies was then counted. Data represent mean + SEM. *p<0.05, students t-test. N>4 cohorts of animals per condition. (D-F) Relative levels of free glucose (D), glycogen (E), or trehalose (F), in adult control (*w*^*1118*^) and *foxo* mutant *(foxoΔ*^*94*^) flies exposed to normoxia or 1% O_2_ hypoxia for 16 hours. n=15. Data represents mean + SEM. *p<0.05, students t-test. (G) *Ldh* mRNA levels measured by qRT-PCR in control (*w*^*1118*^) and *foxo* mutant (*foxoΔ*^*94*^) following 16 hours of 1% O_2_ hypoxia in adults. Data represent mean + SEM. * p,0.05, two-way ANOVA followed by post-hoc t-test. N>10.

Cells, tissues and organisms adapt to low oxygen by altering their metabolism (Semenza, 2011). In particular, a key adaptation is the upregulation of glycolysis. We therefore checked whether FOXO might be important for controlling glucose metabolism in hypoxic animals. We first measured total glucose levels in adult animals exposed to hypoxia. Control animals exhibited a decrease in glucose levels after 16 hours of hypoxia (Figure 2D). *foxo* mutant flies had lower levels of total glucose in normoxia and these levels were even further depleted upon exposure to hypoxia (Figure 2D). We saw a similar pattern of effects when we measured levels of glycogen (the stored form of glucose) and trehalose (the circulating form of glucose in *Drosophila*). Thus, *foxo* mutants showed a significantly greater decrease in both glycogen and trehalose in hypoxia compared to control animals (Figure 2E, F).

Finally, we investigated expression of lactate dehydrogenase (*ldh)* - a key glycolytic enzyme - in *w*^*1118*^ and *foxoΔ*^*94*^ adult females. We saw that control animals increased their *ldh* mRNA when exposed to hypoxia as has been reported before (Lavista-Llanos et al., 2002; Li et al., 2013) and which is consistent with an upregulation of glycolysis. In contrast, *foxo* mutant animals had increased *ldh* levels in normoxia, and this expression increased significantly further in hypoxia (Figure 2G). Taken together, these data indicated that *foxo* mutants show deregulated control over normal glucose metabolism in hypoxia - they show overproduction of *ldh* and they exhibit a larger depletion of both stored and circulating glucose in hypoxia compared to control animals.

### Hypoxia induces FOXO by inhibiting PI 3 K/ Akt signalling

We next examined how hypoxia induces FOXO. The best-studied cellular response to hypoxia involves induction of the HIF1α transcription factor (called *sima* in *Drosophila*). HIF1α induces expression of metabolic and regulatory genes required for hypoxia adaptation, and in both *Drosophila* and *C elegans*, HIF1α is required for organismal tolerance to low oxygen (Centanin et al., 2005; Jiang et al., 2001). However, we found that FOXO was still relocalized to the nucleus in fat body cells from *sima* mutant larvae exposed to hypoxia (Fig 3A). This suggests that induction of FOXO is independent of the classic HIF1α response.

**Figure 3.**
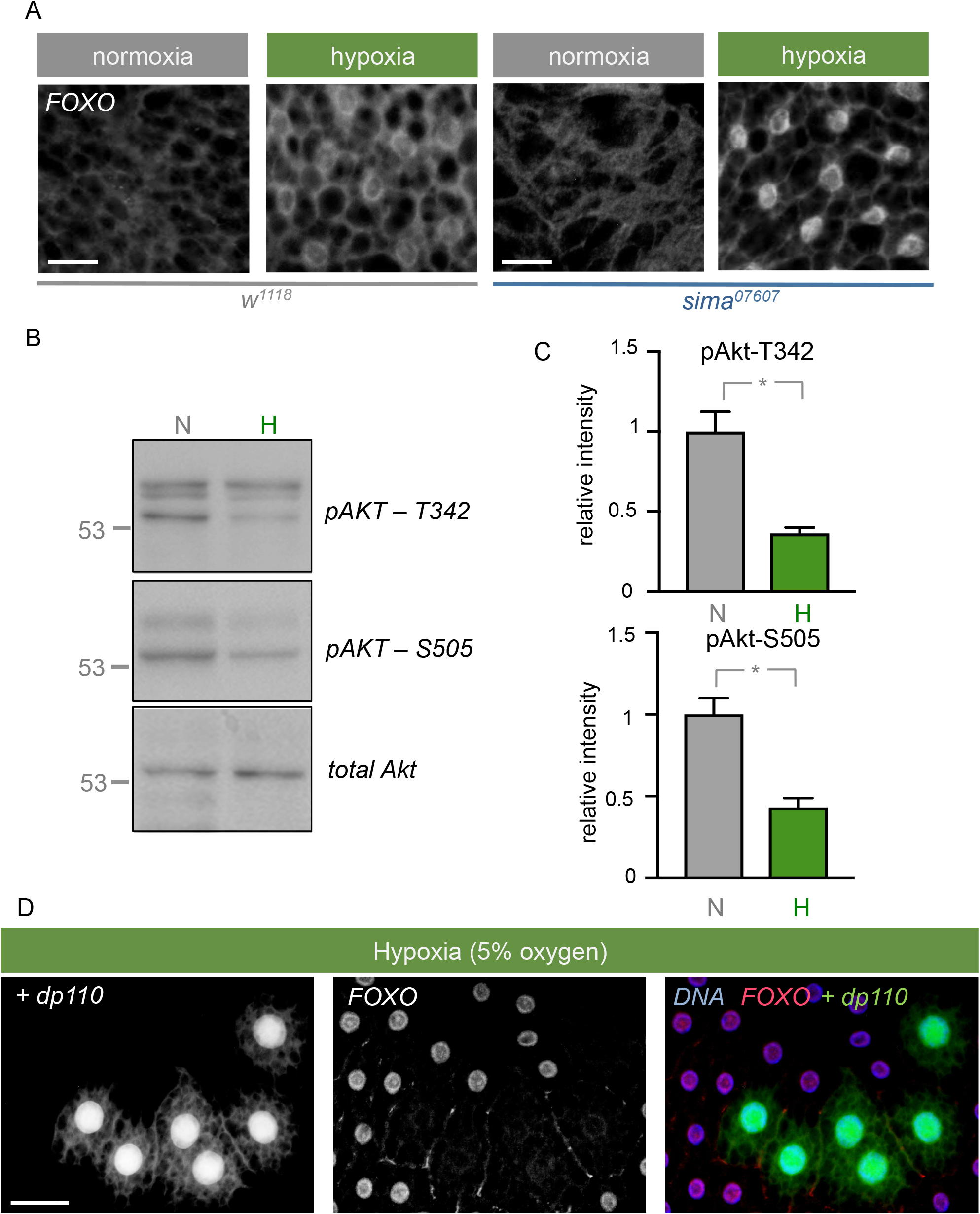
Hypoxia induces FOXO by inhibiting PI3K/Akt. (A) FOXO staining in fat bodies of control (*w*^*1118*^) and *sima* mutant (*sima*^*07607*^) larvae exposed to either normoxia or 5% O_2_ hypoxia for 2 hours. Scale bar is 25 *μ*m. (B,C) Western blot analysis of phosphorylated T342 and S505 Akt, and total Akt in control (*w*^*1118*^) larvae following 2 hours of normoxia (N) or 5% O_2_ hypoxia. Quantification of blots (relative phospho-Akt intensity/total Akt intensity) is shown in (D). N=4 per condition. *p<0.05, students t-test. (D) FOXO staining in UAS-*dp110* overexpressing fat body clones (GFP positive). Nuclei are stained with Hoechst dye (blue). Scale bar is 50 *μ*m.

One main way that FOXO can be regulated is via the conserved insulin/PI3K/Akt pathway (Webb and Brunet, 2014). This is best seen in response to nutrient availability in *Drosophila*. In rich nutrients, insulin signalling to Akt kinase is high and Akt can phosphorylate FOXO, leading its cytoplasmic retention. However, during starvation, insulin/Akt signalling is low, thus reducing phosphorylation of FOXO and allowing it to relocalize to the nucleus to induce transcription. We investigated whether decreased Akt activation was involved in FOXO induction during hypoxia exposure. Akt is activated by phosphorylation at two sites: threonine-342 and serine-505. We measured the relative amounts of Akt phosphorylated at each site after exposure to hypoxia using phospho-specific antibodies. We saw that when third instar larvae were exposed to hypoxia there was a reduction in phosphorylation of Akt at both sites (Figure 3B, C). To determine if suppression of Akt signalling was mediating the induction of FOXO, we used the flp-out technique to induce mosaic expression of the catalytic subunit of PI3K, *dp110*, to maintain Akt activity in fat body cells. We found that during hypoxia, expression of *dp110* was sufficient to prevent FOXO nuclear relocalization (Figure 3D). Taken together, these data show that FOXO induction is mediated by hypoxia-induced suppression of Akt signalling.

### FOXO induces Relish-dependent hypoxia survival

In *Drosophila*, FOXO maintains tissue and organismal homeostasis in response to various stresses, including starvation, oxidative stress, irradiation, and infection. In each case, FOXO functions by regulating diverse and often distinct target genes. We surveyed potential FOXO targets that might be important for hypoxia tolerance and we identified a role for the NF-κB transcription factor *relish*.

In *Drosophila* there are three NF-κB transcription factors, Relish, Dorsal and Dif. They have been best characterized as effectors of immune signalling downstream of the IMD (Relish) and Toll (Dorsal and Dif) pathways, where they induce expression of antimicrobial peptides and promote innate immune responses (Buchon et al., 2014). We found that when exposed to hypoxia, adult *Drosophila* showed an increase in *relish* as reported previously (Bandarra et al., 2014; Liu et al., 2006), but not *dorsal* or *dif*, mRNA levels (Figure 4A-C). Furthermore, we found that this hypoxia-induced increase in relish was blocked in both *foxo* mutant adults (Figure 4D) and larvae (Figure S3). Finally, we found that hypoxia could induce strong expression of Relish-regulated antimicrobial peptides in both adults (Figure E, F) and larvae (Figure S3) and that this was also blocked in *foxo* mutants. These data suggest that in hypoxia, FOXO can induce an immune-like response via upregulation of Relish. To test whether this immune-like response was important for hypoxia survival, we examined hypoxia survival in two independent *relish* null mutants, *rel*^*E38*^ and *rel*^*E20*^ (Hedengren et al., 1999). We found that both *rel*^*E38*^ and *rel*^*E20*^ adult flies showed a significant decrease in viability after hypoxia exposure (Figure 4G, H). Together, these data point to FOXO activation as a meditator of hypoxia tolerance via induction of an immune-like response through the NFκB-like transcription factor *relish*.

**Figure 4.**
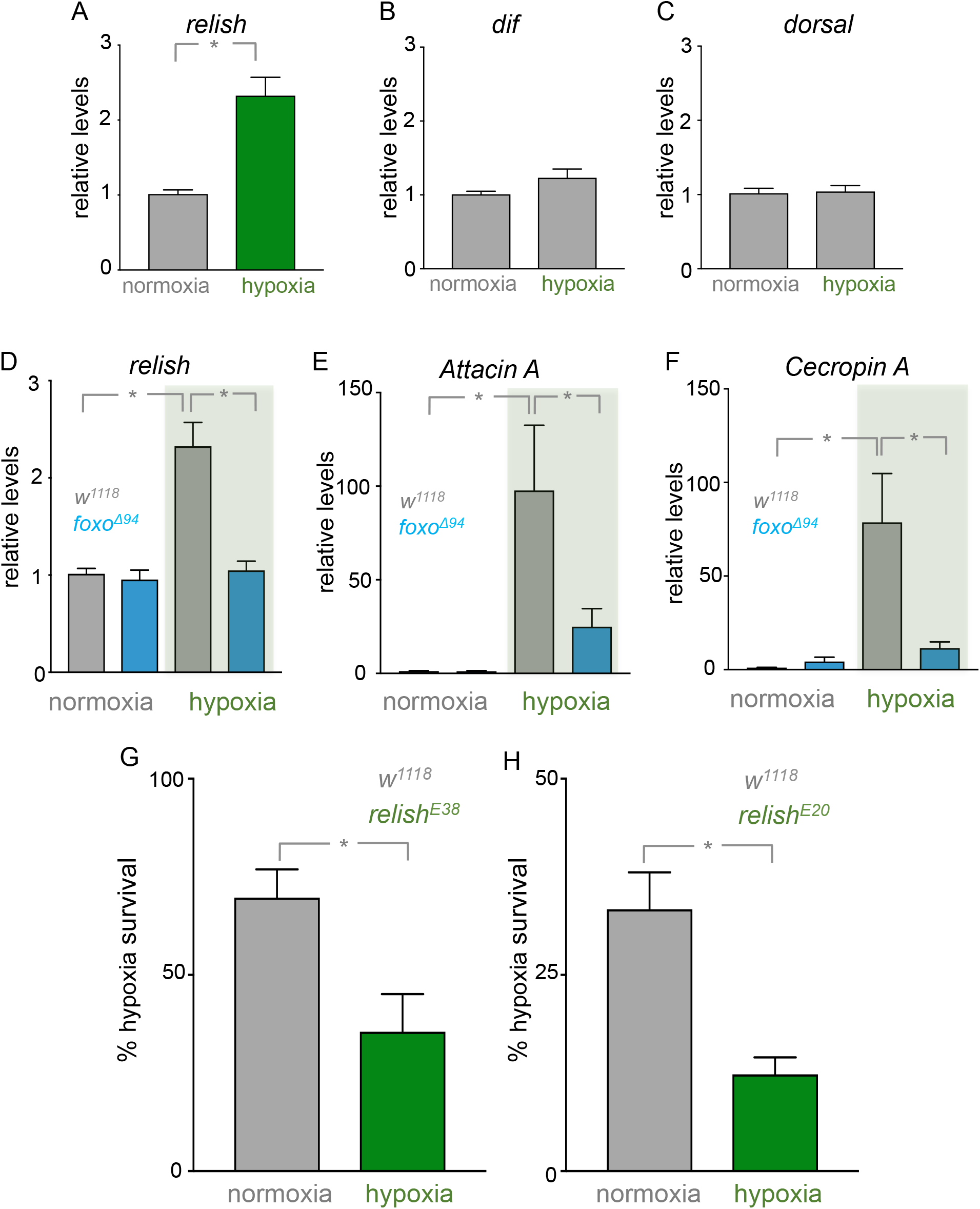
FOXO induces Relish-dependent hypoxia survival. (A-C) Expression levels of *relish* (A), *dif* (B), and *dorsal* (C) mRNA in *w*^*1118*^ adult females exposed to either normoxia or 16 hours of 1% O_2_. Data represent mean + SEM, N=10, *p<0.05, students t-test. (D-F) Expression levels of *relish* (D), *attacin A* (E), and *cecropin A* (F) mRNA in *w*^*1118*^ and *foxoΔ*^*94*^ adult females exposed to either normoxia or 16 hours of 1% O_2_. Data represent mean + SEM, N=10, *p<0.05, 2-way ANOVA followed by students t-test. (G, H) Survival of adult female *w*^*1118*^ or (G) *relish*^*E38*^ or (H) *relish*^*E20*^ flies after exposure to 24 hours of 1% O_2_. Data represents mean + SEM, N= *p<0.05, students t-test.

## DISCUSSION

In this paper, we report that FOXO is a hypoxia-inducible factor required for organismal survival in low oxygen. We saw that this induction of FOXO occurs via suppression of PI3K/Akt signalling. This response is most likely induced by hypoxia-mediated reduction of insulin release and signalling – the main activator of PI3K/Akt - as previously reported in *Drosophila* larvae (Texada et al., 2019; Wong et al., 2014). Interestingly we found that the induction of FOXO upon hypoxia occurs in *sima* mutants suggesting that the FOXO hypoxic response occurs independently of the classically described HIF-1α response. Reduced insulin signalling and FOXO induction have been shown to confer hypoxia tolerance in *C elegans* (Mendenhall et al., 2006; Menuz et al., 2009; Scott et al., 2002). Moreover, the mammalian FOXO homolog FOXO3a can be induced in cell culture upon hypoxia exposure, where it regulates metabolic responses and cell death (Bakker et al., 2007; Jensen et al., 2011). Thus, the induction of FOXO is likely to be a conserved mechanism of hypoxia tolerance in animals.

A central finding of our work is that one way that FOXO provides protection in low oxygen is through induction of an immune-like response. In *Drosophila,* there are two main immune effector pathways that respond to pathogen infection and that work through induction of NF-κB transcription factors – the IMD pathway which targets the NF-κB homolog, Relish, and the Toll pathway which works via the Dorsal and Dif NF-κB transcription factors (Buchon et al., 2014). We found that hypoxia specifically induced Relish via FOXO, and that this response was required for hypoxia tolerance. These data, together with previous work showing hypoxia induction of Relish (Bandarra et al., 2014; Liu et al., 2006), suggest that induction of an immune-like response may be a protective mechanism in low oxygen in *Drosophila*. In the context of animal immunity, there is increasing appreciation of the role for infection tolerance as a defense strategy against pathogens (Ayres and Schneider, 2012; Lissner and Schneider, 2018; Medzhitov et al., 2012). This tolerance is often mediated via alterations in systemic metabolism and physiology to limit infection-induced tissue damage (Ganeshan et al., 2019; Wang et al., 2016; Weis et al., 2017). Our findings suggest that tolerance to hypoxia may share some of these immune functions. In *Drosophila*, this interplay between hypoxia and innate immune responses may reflect the natural ecology of flies. In the wild, *Drosophila* grow on rotting, fermenting food, an environment rich in microorganisms, including pathogenic bacteria. In these anaerobic conditions, low ambient oxygen may ‘prime’ animals to deal with subsequent pathogenic bacterial encounters. Hence, one speculative idea is that experimental exposure of *Drosophila* to hypoxia may induce Relish and provide protection against the detrimental effects of subsequent pathogenic infection. This concept of hypoxia preconditioning has been observed in *C elegans* where it is important in protecting against cell death and damage induced by pore-forming toxins (Bellier et al., 2009; Dasgupta et al., 2007).

Functional interactions between FOXO and Relish have been described in response to other stressors in *Drosophila*. For example, nutrient starvation induces Relish in larvae via FOXO and this is important for controlling systemic insulin signalling (Karpac et al., 2011). In addition, as adults age, FOXO is induced in the intestine and it, in turn, upregulates Relish to control intestinal homeostasis and lifespan (Guo et al., 2014; Karpac et al., 2013). Interestingly, Relish and FOXO have an antagonistic relationship in adult fat and these interactions are important for metabolic adaption and survival upon starvation (Molaei et al., 2019). Hence the links between FOXO and relish are likely to be tissue specific, but they may have evolved to function as a general mediator of stress response. Functional links between NF-κB and FOXO have also been reported in mammalian cells (Lin et al., 2004; Thompson et al., 2015), and together with the reported induction of NF-κB in hypoxia in mammalian cell culture (Fitzpatrick et al., 2011; Rius et al., 2008), they suggest that the hypoxia-FOXO-NF-κB regulation that we see in *Drosophila* may operate in mammalian cells too.

One key way that cells, tissues and organisms adapt to low oxygen is by altering their glucose metabolism in order to maintain homeostasis (Nakazawa et al., 2016; Xie and Simon, 2017). Our data suggest that one reason that *foxo* mutants may show reduced hypoxia tolerance is that they have deregulated control over glucose metabolism. Thus, we saw that *foxo* mutant animals had low levels of glucose in normoxia and that both stored and circulating forms of glucose were significantly decreased under hypoxia compared to controls. These results suggest FOXO is needed for either gluconeogenesis during stress, as has been reported in *C elegans* (Hibshman et al., 2017), or for proper control of glycolysis. Indeed, we saw that expression of *ldh* is markedly increased in *foxo* mutants. Ldh is a rate-limiting enzyme involved in conversion of pyruvate to lactate, which is a key metabolic event that can drive increased glycolysis, and *ldh* levels have been shown to increase in larvae upon hypoxia exposure (Li et al., 2013). Thus, one possibility is that *foxo* mutant animals may engage in abnormally high levels of glycolysis leading to depletion of glucose and reduced hypoxia tolerance. This is consistent with previous studies in *Drosophila* showing a major role for FOXO as a regulator of metabolic homeostasis in the context of other stress responses such as starvation and pathogenic infection (Dionne et al., 2006; Teleman et al., 2008). For example, FOXO often functions in a tissue specific manner to control systemic sugar and lipid metabolism (Borch Jensen et al., 2017; Karpac et al., 2013; Molaei et al., 2019; Wang et al., 2011; Zhao and Karpac, 2017). These effects have been shown to be important for FOXO to extend lifespan and to promote increased tolerance to stress.

It is possible that the effects of FOXO on metabolism in hypoxia could be mediated via Relish. For example, a recent report showed that Relish was required to control metabolic responses to nutrient deprivation in *Drosophila* (Molaei et al., 2019). Furthermore, constitutive activation of IMD signalling – which signals via Relish – was shown to lead to decreased circulating sugars in adult *Drosophila (Davoodi et al., 2019)*. In mammals, NF-κB is activated in response to cytokines and it functions as a central regulator of immune and inflammatory responses (Zhang et al., 2017). Several studies have shown that an important way that NF-κB works to mediate these effects is through the control of glycolysis and mitochondrial metabolic activity (Mauro et al., 2011; Tornatore et al., 2012). Indeed, links between immunity and metabolism are emerging as important components of infection tolerance in animals (Ayres and Schneider, 2012). Our data suggest the possibility that organisms may also co-opt some of these immune-metabolism interactions to tolerate low oxygen.

## METHODS AND MATERIALS

### *Drosophila* stocks

Flies were raised on standard medium containing 150 g agar, 1600 g cornmeal, 770 g Torula yeast, 675 g sucrose, 2340 g D-glucose, 240 ml acid mixture (propionic acid/phosphoric acid) per 34 L water and maintained at 25°C, unless otherwise indicated. The following fly stocks were used: *w*^*1118*^, *sima*^*07607*^/*TM3,Ser,GFP* (Centanin et al., 2008), *foxoΔ*^*94*^/*TM3,Ser* (Slack et al., 2011), *Thor-LacZ* (Bernal and Kimbrell, 2000), *hsflp; UAS-dp110, act>CD2>Gal4,UAS-*GFP (Britton et al., 2002), *Relish*^*E20*^(Hedengren et al., 1999), *Relish*^*E38*^(Hedengren et al., 1999).

### Hypoxia exposure

For all hypoxia experiments vials containing *Drosophila* were placed into an airtight glass chamber into which a premix of 5%oxygen/95% nitrogen, 1%oxygen/99%nitrogen or 100% nitrogen continually flowed. Flow rate was controlled using an Aalborg model P gas flow meter. Alternatively, for some experiments *Drosophila* vials were placed into a Coy Laboratory Products in vitro O_2_ chamber that was maintained at fixed oxygen levels of 1% or 5% by injection of nitrogen gas.

### Immunofluorescence staining

Larvae were inverted using fine forceps in 1x PBS. Inverted larvae were fixed in 8% paraformaldehyde for 30 minutes, washed in 1x PBS/0.1% TritonX-100 (PBST), and blocked for 2 hours at room temperature in 1x PBS/0.1%Tween20/1% bovine serum albumin. Larvae were then incubated overnight with primary antibody diluted in PAT at 4°C, washed 3 times with 1x PBS with 3% TritonX-100 (PBT) and 2% fetal bovine serum (FBS), and incubated with secondary antibody diluted 1∶4000 in PBT with FBS for 2 hours at room temperature. Larvae were washed with PBT and stained with 1:10000 Hoechst dye for 5 minutes, then washed 3 times more with PBT. Larval tissues were isolated using fine forceps and then mounted on glass slides with cover slips using Vectashield mounting media (Vector Laboratories Inc., CA). The rabbit anti-FOXO antibody was used at 1:500 dilution (a gift from Marc Tatar). Alexa Fluor 568 (Invitrogen) was used as the secondary antibody. Hoechst 33342 (Invitrogen) was used to stain nuclei.

### Quantitative PCR

Total RNA was extracted using TRIzol according to manufacturer’s instructions (Invitrogen; 15596–018). RNA samples were then subjected to DNase treatment according to manufacturer’s instructions (Ambion; 2238 G) and reverse transcribed using Superscript II (Invitrogen; 100004925). The generated cDNA was used as a template to perform qRT–PCRs (ABI 7500 real time PCR system using SyBr Green PCR mix) using specific primer pairs. PCR data were normalized to beta-tubulin levels. Each experiment was independently repeated a minimum of three times. The following primers were used:

beta-tubulin: Forward 5’ ATCATCACACACGGACAGG; Reverse 5’ GAGCTGGATGATGGGGAGTA
4e-bp: Forward 5’ GCTAAGATGTCCGCTTCACC; Reverse: 5’ CCTCCAGGAGTGGTGGAGTA
relish: Forward 5’ TCCTTAATGGAGTGCCAACC; Reverse 5’ TGCCATGTGGAGTGCATTAT
dorsal: Forward 5’ TGTTCAAATCGCGGGCGTCGA; Reverse 5’ TCGGACACCTTCGAGCTCCAGAA
dif: Forward 5’ CGGACGTGAAGCGCCGACTTG; Reverse 5’ CAGCCGCCTGTTTAGAGCGG
attacin A: Forward 5’ AGGAGGCCCATGCCAATTTA; Reverse 5’ CATTCCGCTGGAACTCGAAA
cecropin A: Forward 5’ TCTTCGTTTTCGTCGCTCTCA; Reverse 5’ ATTCCCAGTCCCTGGATTGTG

### Lac Z staining

Larvae were inverted using fine forceps in 1x PBS. Inverted larvae were fixed in 8% paraformaldehyde for 30 minutes, washed in 1x PBS-0.1% TritonX-100 (PBST), and then incubated in 500μl of an X-Gal solution containing10 mM sodium phosphate buffer, pH 7.2, 150 mM NaCl, 1mM MgCl_2_, 10 mM K_4_[Fe^II^(CN)_6_], 10 mM K_3_[Fe^III^(CN)_6_], 0.1% Triton X-100 with 12.5μl of an 8% X-Gal solution (in DMSO) added immediately prior to incubation. Samples were then incubated at 37C until the X-Gal staining was visible.

### Measurement of hypoxia survival

#### Larvae

newly hatched larvae were placed in food vials (50 larvae per vial) and then maintained in either normoxia or hypoxia (5% oxygen). Larvae exposed to hypoxia were maintained in this environment until about 80% of larvae had pupated. Then, vials were removed from hypoxia and the numbers of eclosing adults were counted.

#### Adults

4-5 days post-eclosion, mated female adults were placed in placed into hypoxia (1% oxygen) for 24 hours in cohorts of 20 flies per vial. Then, vials were removed from hypoxia and the flies were allowed to recover for 48 hours before the number of dead flies were counted.

#### Starvation

At 4-5 days post-eclosion, mated female adults were subjected to starvation by transferring them from food vials to vials containing 0.4% agar/PBS for 24 hours. The number of dead flies was then counted.

### Glucose, glycogen, trehalose and TAG assays

Adult female *Drosophila* were either exposed to hypoxia (1% oxygen) for 16 hours or maintained in normoxia and then frozen on dry ice. Colorimetric assays for each of the metabolites were then conducted using the methods described in detail in (Tennessen et al., 2014).

### Preparation of protein extracts and western blotting

*Drosophila* larvae were lysed with a buffer containing 20 mM Tris-HCl (pH 8.0), 137 mM NaCl, 1 mM EDTA, 25 % glycerol, 1% NP-40 and with following inhibitors 50 mM NaF, 1 mM PMSF, 1 mM DTT, 5 mM sodium ortho vanadate (Na_3_VO_4_) and Protease Inhibitor cocktail (Roche Cat. No. 04693124001) and Phosphatase inhibitor (Roche Cat. No. 04906845001), according to the manufacturer instructions. Protein concentrations were measured using the Bio-Rad Dc Protein Assay kit II (5000112). Protein lysates (15 μg to 30μg) were resolved by SDS–PAGE and electro transferred to a nitrocellulose membrane, subjected to Western blot analysis with specific antibodies, and visualized by chemiluminescence (enhanced ECL solution (Perkin Elmer)). Primary antibodies used in this study were: anti-Akt (Cell Signaling #9272, 1:500 dilution), anti-pAkt-T342 (gift from Michelle Bland), anti-pAkt-S505 (Cell Signaling #4054, 1:1000 dilution). Secondary antibodies were purchased from SantaCruz Biotechnology (sc-2030, 2005, 2020). For experiments looking at Akt phosphorylation, total Akt levels were used as a loading control because the level of this protein was unaffected by hypoxia.

### Statistical analyses

Data were analyzed by Students t-test or two-way ANOVA. All statistical analysis and data plots were performed using Prism software. In all figures, statistically significant differences are presented as: * and indicate p<0.05.

## ACKNOWLEDGEMENTS

We thank Edan Foley, Linda Partridge, Bruce Edgar, Mark Tatar for the gift of reagents and fly stocks. Stocks obtained from the Bloomington *Drosophila* Stock Center (NIH P40OD018537) were used in this study. This work was supported by a NSERC Discovery grant to S.S.G. E.C.B was supported by an Alberta Innovates Health Solutions Graduate Studentship. A.N.B-P was supported by an NSERC summer studentship. D.M.P was supported by an NSERC CGS-M graduate scholarship.

---

**Figure S1.**
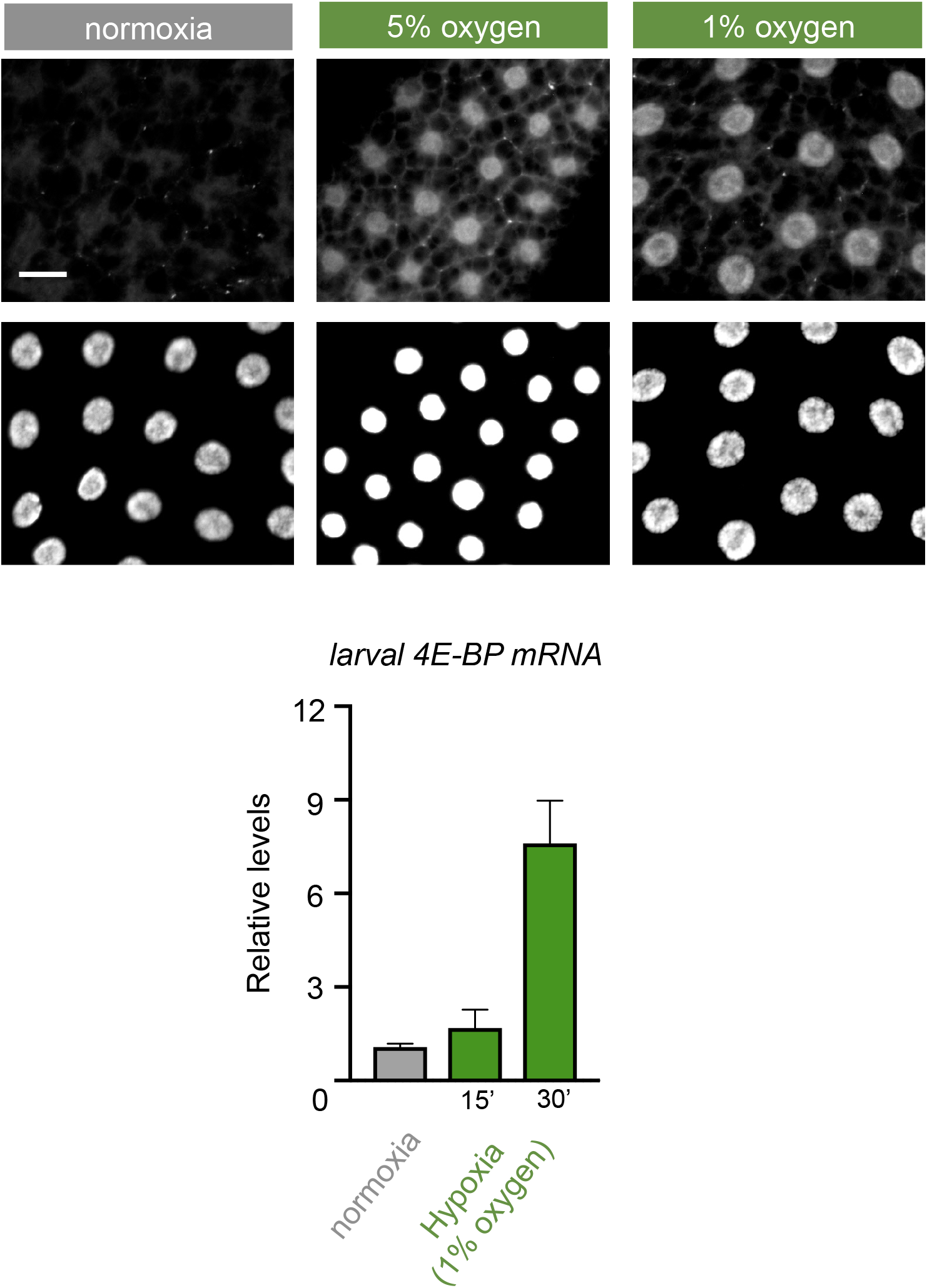
FOXO is induced rapidly in hypoxia. (A) FOXO staining of 96-hour AEL *w*^*1118*^ larval fat bodies following exposure to hypoxia for 15 minutes. Nuclei are stained with Hoechst (bottom panels). Scale bar is 25 *μ*m. (B) *4e-bp* mRNA levels measured by qRT-PCR in control (*w*^*1118*^) larvae exposed to either normoxia or hypoxia (1% oxygen) for 15 or 30 minutes. Data represent mean + SEM, N=10, *p<0.05, students t-test.

**Figure S2.**
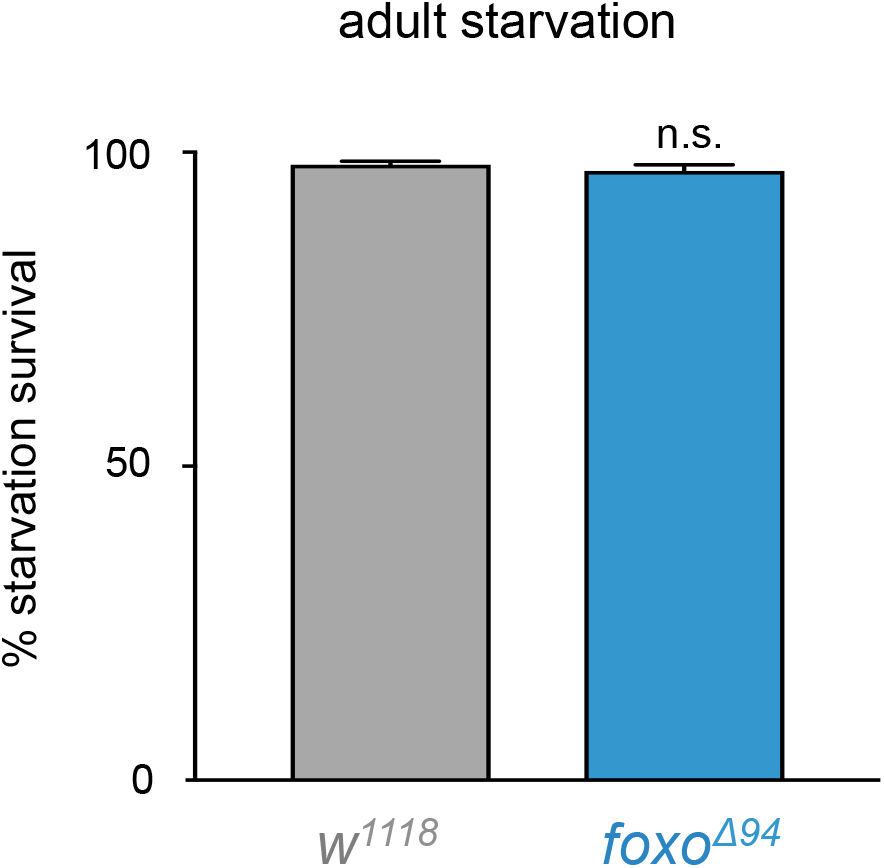
*foxo* mutant survival is not affected by short term nutrient deprivation. (A) Survival of adult female *w*^*1118*^ and *foxoΔ*^*94*^ flies 2 days after starvation for 24 hours. Data represented as mean + SEM for n=4 groups of 20 flies.

**Figure S3.**
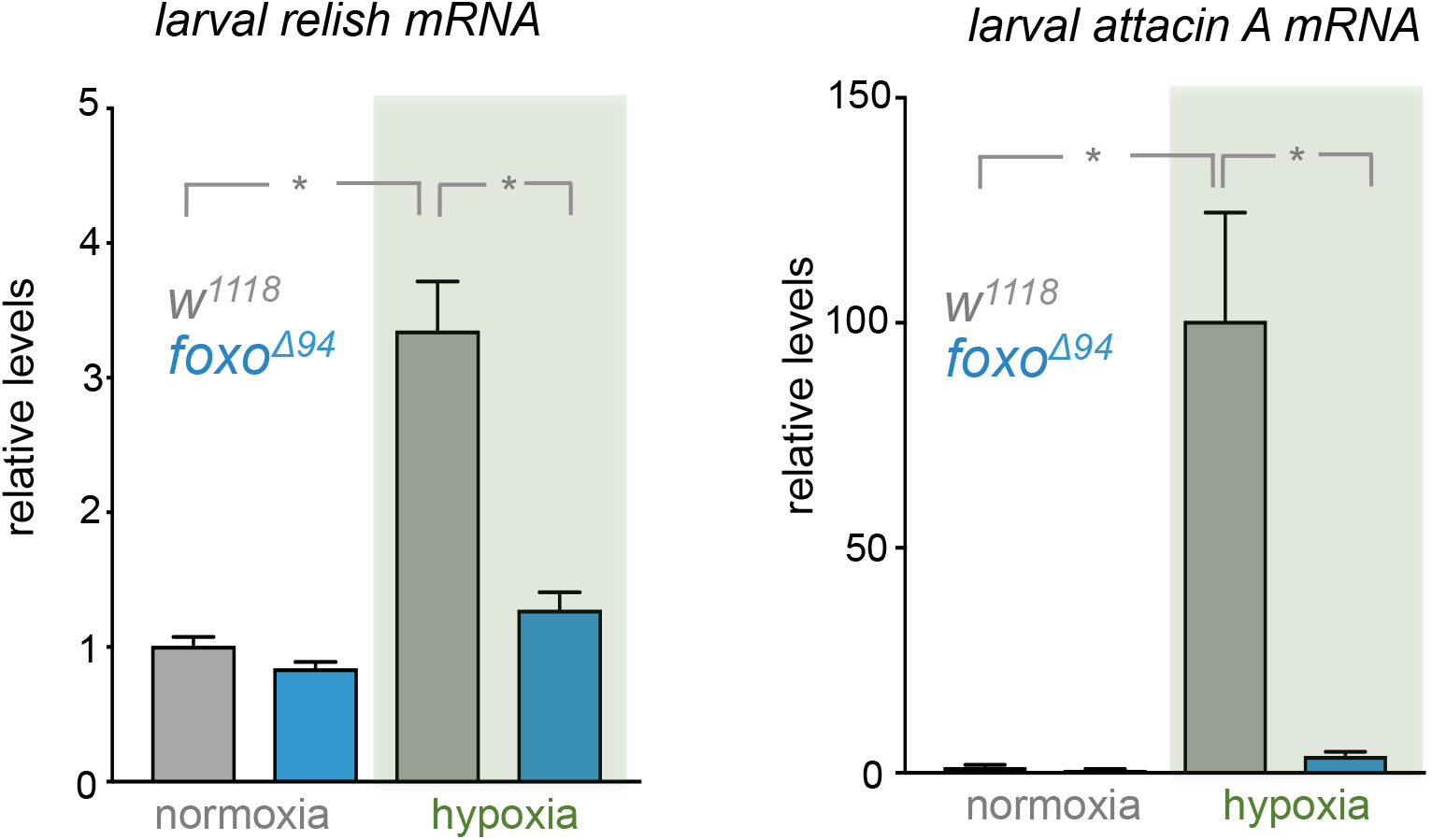
*relish* is induced by FOXO in hypoxic larvae. Expression levels of (A) *relish* or (B) *attacin* A mRNA in *w*^*1118*^ and *foxoΔ*^*94*^ larvae exposed to 5% O_2_ for 6 hours. Data represent mean + SEM, N=10, *p<0.05, 2-way ANOVA followed by students t-test.

